# Inference of Population Admixture Network from Local Gene Genealogies: a Coalescent-based Maximum Likelihood Approach

**DOI:** 10.1101/2020.05.04.076075

**Authors:** Yufeng Wu

**Affiliations:** Department of Computer Science and Engineering, University of Connecticut, Storrs, CT 06269, U.S.A.

**Keywords:** Population admixture, population genetics, coalescent theory, multispecies coalescent, maximum likelihood inference

## Abstract

Population admixture is an important subject in population genetics. Inferring population demographic history with admixture under the so-called admixture network model from population genetic data is an established problem in genetics. Existing admixture network inference approaches work with single genetic variation sites. While these methods are usually very fast, they don’t fully utilize the information (e.g., linkage disequilibrium or LD) contained in population genetic data. In this paper, we develop a new admixture network inference method called GTmix. Different from existing methods, GTmix works with local gene genealogies that can be inferred from population haplotypes. Local gene genealogies represent the evolutionary history of sampled alleles and contain the LD information. GTmix performs coalescent-based maximum likelihood inference of admixture networks with the inferred genealogies based on the well-known multispecies coalescent (MSC) model. GTmix utilizes various techniques to speed up likelihood computation on the MSC model and optimal network search. Our simulations show that GTmix can infer more accurate admixture networks with much smaller data than existing methods, even when these existing methods are run with much larger data. GTmix is reasonably efficient and can analyze genetic datasets of current interests.

## Introduction

Population demographic history is a complex interplay of various processes, such as population divergence and isolation, population size changes, migration and admixture. The simplest population demographic history model is the population tree model (similar to phylogenetic tree for species evolution on the high level), which only models population divergence. In practice, however, the population tree model is often too simplistic for most population genetic study. One key missing aspect by the population tree model is gene flow. In this paper, we focus on population admixture, which is one of the most important types of gene flow. Population admixture is often so widespread that admixture has to be addressed in most demographic history studies. Human populations, for example, are known to be strongly affected by admixture throughout the human history [21, 18]. When admixture is considered, population history becomes a network (called admixture network). In addition to modeling population divergence, admixture network has admixture nodes that model admixture events. See Figure 1(a) for an illustration of admixture network.

**Figure 1:**
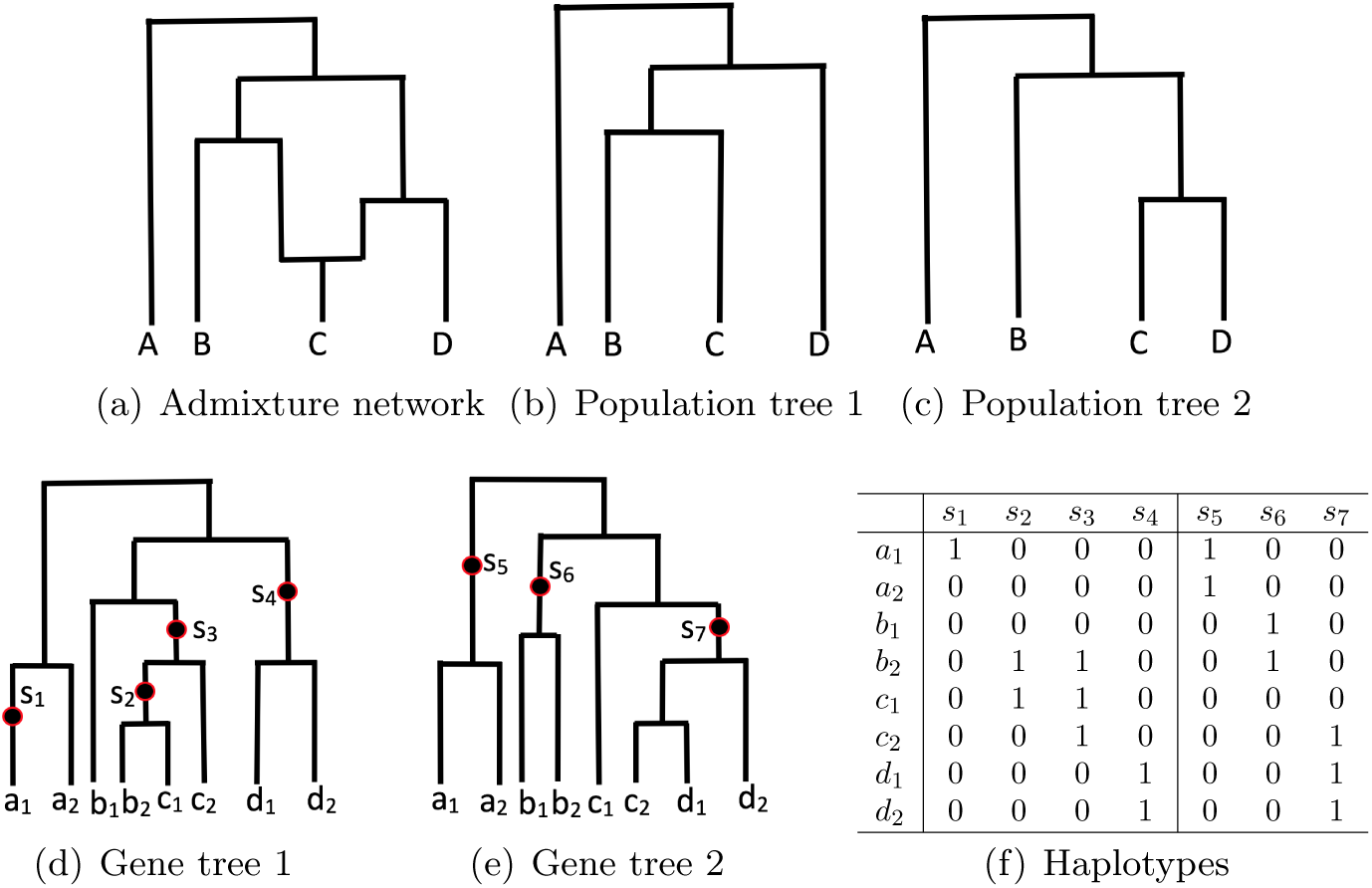
Illustration of admixture network. Network is shown in part 1(a). Four populations. C is admixed. Two population trees in this network are shown in parts 1(b) and 1(c). Parts 1(d) and 1(e): two gene trees at different loci. Two sampled alleles per population: alleles a_1_ and a_2_ are from the population A, alleles b_1_ and b_2_ are from the population B and so on. Darkened dots: mutations. Mutations: follow infinite sites model. Part 1(f): haplotypes for the two gene trees.

Since demographic history is not directly observable, it is highly desirable to infer population history with admixture (i.e., admixture network) from extant population genetic data [20, 13]. Inferring admixture networks from population genetic data is challenging computationally. First, the space of admixture networks can be very large even for moderate number (say ten) of populations and small number (say two) of admixture events^1^. Moreover, the effect of admixture on population genetic data is subtle and is not easily detectable. Meiotic recombination further complicates the situation by breaking the linkage between genetic variation sites.

A natural population genetic model for admixture network inference is the coalescent model [11]. Coalescent process determines stochastically how sampled alleles coalesce in a population. Since there are multiple populations in an admixture network, the underlying coalescent is the multispecies coalescent (MSC). Multispecies coalescent is the fundamental genetic model for the study of multiple populations (species) [23]. Multispecies coalescent can be extended to allow gene flow (see, e.g. [32]). Under the multispecies coalescent model, one can (at least in principle) perform likelihood-based inference of admixture networks. However, inference based on multispecies coalescent is computationally intensive [23, 4]. Due to these difficulties, existing methods for inferring admixture networks usually perform inference with allele frequencies at individual genetic variation sites on somewhat simplified genetic models. For example, the TreeMix approach [20] infers admixture networks from allele frequency data based on a Gaussian approximation of genetic drift. MixMapper [13], another method for network inference, has similar high-level approach as TreeMix. Both TreeMix and MixMapper are fast to handle large genetic data. There are several downsides of these existing methods. First, only working with allele frequencies at individual sites may potentially lose information, especially the linkage disequilibrium (LD) among nearby sites. Moreover, these approaches are based on approximations of the underlying genealogical process, which may not very accurately model the MSC process. Our experience indicates that TreeMix and MixMapper, while useful, do not provide very accurate inference results in some of our simulations.

In this paper, we present a new method for inferring admixture networks from population genetic data. Our new method is named GTmix (which stands for <u>G</u>ene <u>T</u>ree based ad<u>mix</u>ture network inference). The following summarizes the main features of GTmix.

1. GTmix works with haplotypes, rather than individual variation sites. This allows GTmix to exploit the LD information from genetic data, which is not considered by TreeMix and MixMapper. GTmix doesn’t directly perform inference from haplotypes. Instead, GTmix takes input in the form a set of gene genealogies 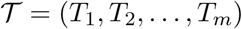 that are *inferred* from the given haplotypes. Here, *T*_*i*_ is the inferred gene genealogy at a site *s*_*i*_, which represents the evolutionary history of haplotypes at *s*_*i*_. Due to recombination, genealogies at different loci may be different. Gene genealogy is arguably more informative than individual genetic variation sites. A traditional view in population genetics is that while gene genealogy is informative, there is little power in inferring gene genealogies from genetic data largely due to recombination (see, e.g., [27]). Recently, however, local genealogy inference is being actively studied and applied in population genetics. There are several existing tools for inferring gene genealogies from population haplotypes [22, 15, 24, 9]. Among these tools, RENT+ [15] is efficient and easy to use. So we use RENT+ for local genealogy inference in this paper.
2. GTmix is a maximum likelihood approach: it aims aITt finding an admixture network 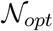 such that 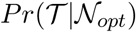 is maximized. Here, 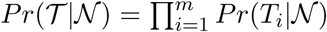 (based on the assumption the independence of each *T*_*i*_). 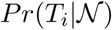 is the probability of observing genealogy *T*_*i*_ for the admixture network 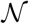 under the multispecies coalescent with admixture (MSCA) model. While in general computing 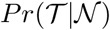 under the MSCA model is computationally challenging, GTmix takes advantage of the recent algorithmic progress in probability computation on the MSC model [29, 31, 19], which allows such probability to be computable in practice for medium size data. By working on the MSCA model, GTmix performs admixture network inference on an arguably more rigorous model than existing methods (such as TreeMix).
3. GTmix can be configured to infer admixture networks with arbitrary number of admixture events, although the running time will increase with the number of admixture events. It can infer recent admixture (i.e., that forms an extant population) and also ancient admixture (i.e., that forms an ancestral population). GTmix is reasonably efficient computationally. In Phylogenetics, there exists methods (most notably the program Phylonet [26]) for inferring phylogenetic networks (somewhat related to admixture networks) based on the MSC model. Our experience indicates that GTmix can perform likelihood based inference on data with significantly larger number of taxa and alleles than that allowed by Phylonet.

Through simulation, we show that while GTmix is not as fast as existing methods such as TreeMix, GTmix is more accurate than existing methods in many simulations. In particular, GTmix can infer more accurate admixture networks under many settings using much smaller amount of data than existing methods, even when those existing methods use much larger data than GTmix. Our results suggest that inferred genealogies can indeed be informative for population demographic history inference. Also coalescent-based probabilistic inference with local genealogies can be potentially more accurate than existing methods using single variation sites.

## 2 Background

### 2.1 Admixture network

Admixture network is conceptually similar to phylogenetic network as studied in the Phylogenetics literature [8, 16]. Admixture is a natural extension to the population tree model (see, e.g., [30]). Admixture network (or simply network) is a directed acyclic graph 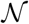, and is leaf-labeled by a set of populations. Moreover,

1. Admixture nodes are those nodes in 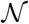 with two incoming edges. The two ancestral nodes of an admixture node are called the source populations.
2. For a network 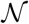, when only *one* of the incoming edges of each admixture node is kept and the other is deleted, we always obtain a tree *T*. This tree is called population tree.

Note that in admixture network, a node refers to a population. An admixed node has two incoming edges, which originate from two source populations for the admixed population. So for an admixture node *v* in a network, there are two ancestral nodes: the left parent (denoted as *Left*(*v*)) and the right parent (denoted as *Right*(*v*)). Throughout this paper, we assume each admixture is two-way (i.e., formed by two ancestral populations). More general admixture (involving three or more ancestral populations) can in principle be converted to a chain of two-way admixture events.

#### Population tree

Recall that a population tree *T* is contained in a network 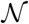 when we remove one of the two incoming edges at each admixture node. Suppose there are *n*_*a*_ admixture nodes in 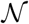. There are 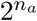 population trees contained in 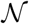. In Figure 1, there are two population trees contained in the network, as shown in Figures 1(d) and 1(e). Note that population trees don’t capture all the possible demographic histories in 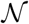. To see this, look at the network in Fig. 1(a) and the two population trees in Figures 1(b) and 1(c). Here, *C* is either sibling of *B* or *D* in a population tree. However, this doesn’t capture the possible scenarios that, for example, one lineage from *C* coalesces with a lineage from *B*, while the other from *C* coalesces with a lineage from *D*. Despite this shortcoming, GTmix relies on population trees in computing the coalescent likelihood because it is computationally more efficient to work with population trees, instead of the network.

#### Branch length

Branches in an admixture network have lengths in the standard coalescent units. This accommodates the case where different branches have different population sizes. In this paper, we assume population sizes remain constant in a branch of 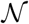.

#### Admixture proportions

At each admixture node *v*, there is an admixture proportion *f*_0_(*v*) where 0 ≤ *f*_0_(*v*) ≤ 1. *f*_0_(*v*) refers to the proportion of alleles of population *v* originating from the left parent *Left*(*v*). We define *f*_1_(*v*) = 1.0 − *f*_0_(*v*) as the proportion of alleles of *v* from the right parent *Right*(*v*). Admixture proportions determine the probability of individual population trees in 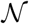.

#### Genealogy and multispecies coalescent

Gene genealogy and multispecies coalescent are central to the GTmix method. Due to the lack of the space, background information on genealogy and multispecies coalescent is given in the Supplemental Materials.

## 3 Method

### 3.1 The high-level approach

In this paper, we develop a new admixture network inference tool, called GTmix. We assume haplotypes *𝓗* from *n*_*p*_ populations are given. We further assume the number of admixture events *n*_*a*_ is known. Our goal is inferring the maximum likelihood admixture network 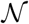 with *n*_*a*_ admixture nodes for these populations from 𝓗. One natural approach is finding 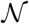 that maximizes the likelihood of 𝓗. However, computing the likelihood 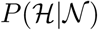 is challenging. GTmix uses the following techniques to develop a practical coalescent-likelihood based inference approach.

1. GTmix works with local genealogies 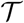 which are inferred from 𝓗, not 𝓗 directly. Before using GTmix, we first use a local genealogy inference tool (e.g., RENT+ [15]) to infer local gene genealogies 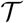 from 𝓗. The main benefit of working with inferred genealogies is that computing the likelihood of 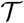 is easier than directly computing the likelihood of 𝓗 for 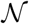; also local genealogies 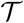, while often noisy, capture the important LD information, where single SNP sites don’t. The number of genealogies can be very large in practice. GTmix uses a filtering scheme to choose more reliable gene genealogies for inference.
2. GTmix takes the inferred 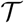 to infer 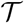 that maximizes 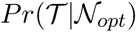. 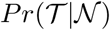 is computed approximately by summing up the probability of 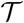 for each population tree in 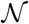. The latter probability is computed using the fast gene tree probability algorithm in [19]. This simplified probability can be computed relatively fast when the network is not too large. GTmix implements effective methods to perform admixture network search.

### 3.2 Inferring local gene genealogies

GTmix takes a list of local gene genealogies 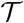 as input. 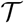 can be inferred by applying a local genealogy inference tool on the given haplotypes 𝓗. GTmix is tested with genealogies inferred by the program RENT+ [15], although other tools (e.g., [24, 9]) can be used as well. Gene genealogies are inferred from haplotypes of each locus separately. RENT+ infers a (rooted binary) genealogy for each SNP site within the locus. The total number of genealogies can be very large because there can be different genealogy at each SNP. This makes GTmix infeasible to perform inference on all genealogies since computing the probability of a genealogy is not trivial. To develop a practical method, GTmix samples a subset of genealogies (trees) using a two-stage approach: (i) it first chooses a subset of (likely more reliable) trees from each locus, and (ii) it then chooses a smaller subset of trees from the list of trees from all loci. Due to the lack of space, the details are given in the Supplemental Materials.

### 3.3 Gene tree probability computation for admixture network

GTmix takes a list of inferred genealogies 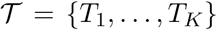 as input. The most important aspect for inferring admixture network 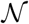 is computing 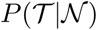 on the multispecies coalescent with admixture (MSCA) model. Assuming the independence of genealogies, 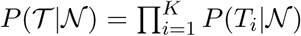. Computing 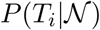 for a network 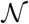 is not trivial (see, e.g., [32]). The main difficulty for computing 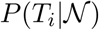 is that the number of feasible coalescent histories can be very large under the MSC model. To develop a practical method that can work with relatively large data, GTmix takes an approximation here: instead of computing the probability of *T*_*i*_ for 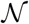, GTmix computes the probabilities of *T*_*i*_ for each population tree in 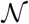 and then use the sum of these probability as an approximation of 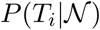. While this is only an approximation, our experience indicates this allows reasonably accurate and efficient inference of admixture network.

To be specific, we consider all possible population trees in 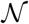. Each population tree is obtained by picking one of the two incoming admixture edges at admixture node *v* with probabilities *m*_*v*_ and 1.0 − *m*_*v*_ respectively. Let the embedded population trees within 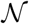 be 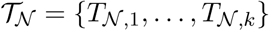. Let *n*_*a*_ be the number of admixture notes in 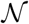. A population tree 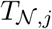 is associated with a binary vector *D*_*j*_ of length *n*_*a*_. *D*_*j*_[*a*] = 0 (respectively *D*_*j*_[*a*] = 1) if the left (respectively right) incoming edge is chosen to obtain 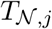. Here, each 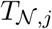 has a probability of being the population tree: 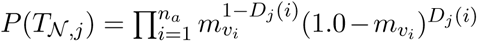. Here, *v*_*i*_ is the *i*-th admixture node. Recall that branches in 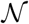 have lengths. So trees in 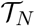 have branch lengths. In contrast, we assume genealogical trees in 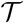 are topologies only. Thus,

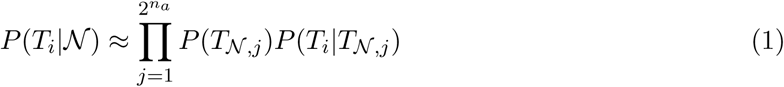

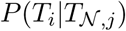 is exactly the gene tree probability of a gene tree topology for a fixed population tree (with branch lengths). GTmix uses the approximate gene tree probability algorithm in [19]. This is because this algorithm is much more scalable than exact gene tree probability algorithms in [4, 29, 31]. Note that Equation 1 is only an approximation. It assumes all lineages at an admixture node are inherited from a single parent. In reality, when there are multiple lineages from an admixed population, different lineages may trace to different parental populations of this population. To see this, we again look at the network in Fig. 1(a) and the gene tree in Fig. 1(d). Suppose at the time of admixture, lineages *c*_1_ and *c*_2_ remain un-coalesced. Then, Equation 1 ignores the situation where, for example, *c*_1_ is from the left and *c*_2_ is from the right. We adopt this approximation because: (i) computing gene tree probability for a population tree is much faster than computing the gene tree probability for a network, and (ii) empirical tests suggest that this approximation appears to provide reasonably accurate inference.

### 3.4 Finding the maximum likelihood admixture network

GTmix assumes the number of admixture events *n*_*a*_ in the network is known. Note that existing network inference methods such as TreeMix and MixMapper also make the same assumption. GTmix finds the maximum likelihood admixture network using a simple iterative procedure:

1. Construct an initial network 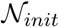 with *n*_*a*_ admixture nodes from the inferred gene genealogies 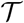. Let 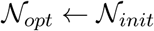.
2. Find the set of admixture networks that are similar topologically to 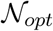 and have *n*_*a*_ admixture nodes.
3. Let 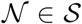 that maximizes the likelihood 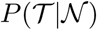. If 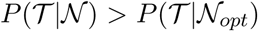, set 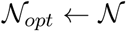, and go to step 2. Otherwise, stop.

#### 3.4.1 Constructing the initial network

GTmix constructs an initial network 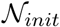 from a set of genealogies 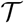 as follows. It first constructs a network 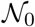 without admixture nodes (i.e., a population tree) by neighbor joining (NJ). Then, GTmix adds *n*_*a*_ admixture nodes to 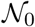, which gives the initial network 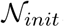. To run the NJ algorithm, GTmix estimates the pairwise distance between each pair of populations *p*_*i*_ and *p*_*j*_ based on topological information in 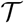. This is shown in Algorithm 2 (in the Supplemental Materials). The pairwise population distance between *p*_*i*_ and *p*_*j*_ is estimated based on the average distance between two sample alleles from *p*_*i*_ and *p*_*j*_ (in terms of the number of edges separating these two alleles on an inferred genealogy).

GTmix now adds *n*_*a*_ admixture nodes to 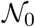 by choosing proper leaf nodes in 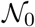 and turning them into admixture nodes. That is, the initial admixture populations are the extant populations in 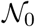. Note that ancestral admixture events (i.e., admixture at ancestral populations) are accommodated during the network search stage. GTmix relies on the so-called minimum deep coalescent (MDC) [17, 14] to identify likely admixed extant populations. Briefly, MDC is a statistic that measures the topological deviation of the given gene trees from a consensus tree based on the MSC model. Refer to [17, 14] for more details on MDC. While MDC is known to have issues in phylogenetics (see, e.g., [5]), we choose to use MDC due to its simplicity and efficiency.

The key idea of inferring likely admixed extant populations by GTmix is that admixed extant populations tend to make gene trees *deviate* significantly from the standard MSC process. This is because the standard MSC process doesn’t consider admixture. Thus, intuitively, an admixed population tends to lead to gene trees that are quite different from gene trees arising from the standard MSC process. Suppose we discard all alleles from the admixed population from the gene trees, the (reduced) gene trees tend to have a better fit to the standard MSC process. GTmix uses the MDC statistic to measure how well gene trees fit the standard MSC process. More specifically, suppose a population *p* is a candidate for admixed extant population. We first compute the MDC score (denoted as *MDC*) of the original 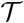. Then we discard all gene lineages from the population *p* in all trees in 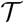. We call the reduced genealogies 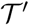. We then compute the MDC score (denoted as *MDC*_*p*_) on 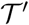. Since each tree in 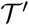 is a subtree of the corresponding tree in 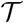, *MDC*_*p*_ ≤ *MDC*. We expect *MDC* − *MDC*_*p*_ for admixed *p* to be significantly larger than *MDC* − *MDC*_*p*0_ for an un-admixed population *p*_0_. This leads to Algorithm 1 for finding likely admixed extant populations.

##### Algorithm 1

Finding likely admixed extant populations from genealogies 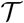

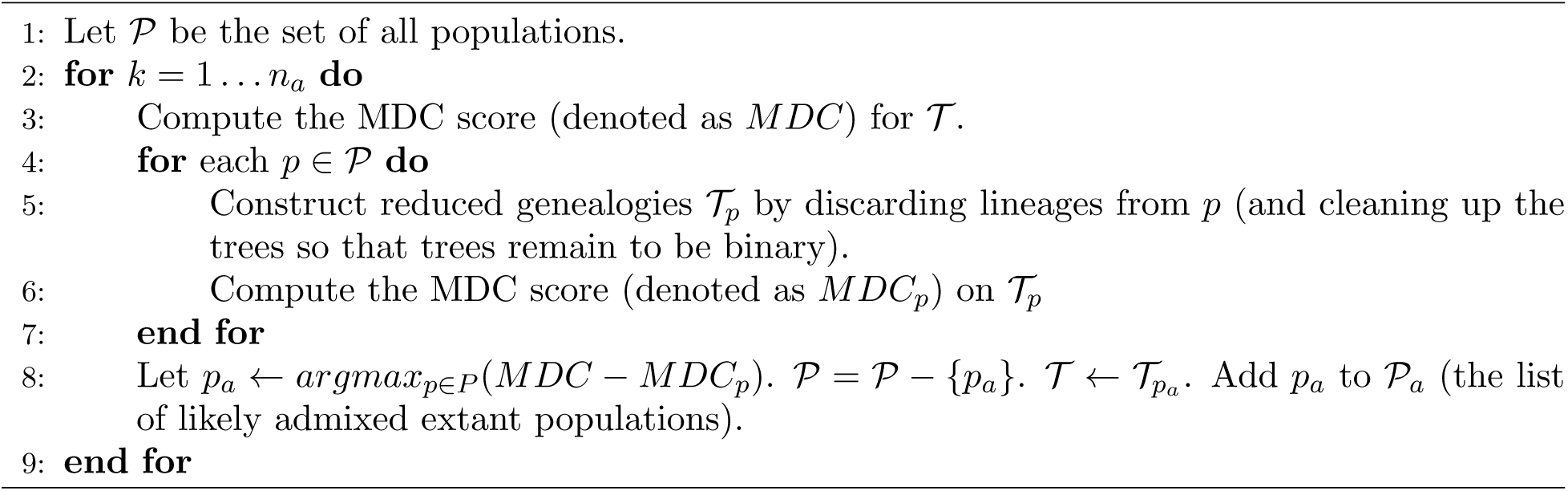

##### Constructing initial network 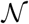 by adding admixture nodes

For each likely admixed extant population 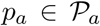, GTmix modifies 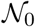 by adding one admixed branch into *p*_*a*_ to make *p*_*a*_ an admixture node. GTmix considers all possible admixture source populations that can be ancestral to *p*_*a*_ according to the current network. Each choice leads to a network 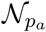. Then it chooses one source population that gives the maximum probability 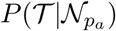. After this step, we obtain the initial network 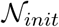.

#### 3.4.2 Search over the space of networks

GTmix searches for the set of topologically similar networks 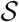 from the current network *N*_*opt*_. Networks in 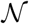 are obtained by applying one nearest neighbor interchange (NNI) to the current network. More specifically, consider each branch *b* = (*v*_*p*_, *v*_*c*_) in 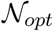, where node *v*_*p*_ is the parent of node *v*_*c*_. Let *v*_*s*_ be the sibling of *v*_*c*_, and *v*_*sp*_ be the sibling of *v*_*p*_. Then after applying one NNI on *b*, we obtain a new network with *v*_*sp*_ being the sibling of *v*_*c*_ and *v*_*s*_ being the sibling of *v*_*p*_. That is, the NNI swaps *v*_*s*_ with *v*_*sp*_ which leads to a new network 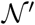. For each network 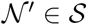, GTmix uses hill climbing to find the network that maximizes the likelihood.

##### Simulation

Due to the lack of space, simulation procedure is given in the Supplemental Materials.

## 4 Results

Due to the lack of space, some results are provided in the Supplemental Materials.

### 4.1 Results on simulated data

At present, GTmix cannot run on data with large number of populations and/or large number of alleles per population. Thus, we run GTmix with relatively small number of alleles per population (by default, four alleles per population). Existing methods such as TreeMix and MixMapper are designed to work with data with relatively large number of alleles. For comparison, we run TreeMix and MixMapper on two settings: (i) the “small” data: the same data as given to GTmix, and (ii) the “large” data: simulated data with much larger number of alleles per population (by default, 100 alleles per population, i.e., 25 times of the small data). Note that the number of combinations of different parameters is very large. In the following, when we evaluate the effect of a particular parameter, we keep all other parameters to be the default values.

#### 4.1.1 Metrics for benchmarking the accuracy of admixture network inference

In this paper, we use the following two metrics for comparing the topology of 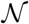 and that of 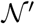.

1. Best-match population tree inference error. Recall that a network 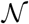 with *n*_*a*_ admixture nodes contains 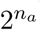 population trees. Assuming 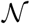 and 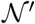 have the same number of admixture nodes, we compare the list of 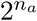 population trees 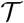 in 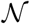 with 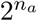 population trees 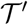 in 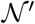. Here we need to match each tree 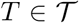 with a tree 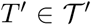. There are 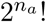 ways of matching 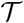 with 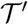. For example, when *n*_*a*_ = 1, there are two ways of matching; when *n*_*a*_ = 2, there are 24 ways of matching. The so-called Robinson-Foulds (RF) distance between a pair of matched trees *T* and *T′* is used. The RF distance is equal to the number of clades (subtrees) that are in *T* but not in *T′*. We normalize the RF distance to be between 0 and 1. For each matching, we take the average RF distance of all matched pairs of trees as the inference error. We take the smallest average RF distance over all matchings as the best-match inference error.
2. Percentage of correctly inferred admixed populations. It can be biologically important to identify which populations are admixed. This is measured by the average percentage of correctly inferred admixed populations.

In addition to topology inference, we also examine the accuracy of the inferred admixture proportions. We use the absolute difference between the inferred admixture proportion and the true admixture proportion as the inference error for admixture proportion. For each settings, we simulate ten replicate datasets. Reported results are the average over these replicates.

#### 4.1.2 Network inference accuracy

##### Varying number of populations

We run GTmix, TreeMix and MixMapper on data with varying number of populations (from 4 to 10 populations). We sample four alleles for each population. We also run TreeMix and MixMapper with large data (100 alleles for each population). Fig. 2 shows the results. We can see that GTmix is significantly better than TreeMix and MixMapper on small data in terms of both the topology inference and admixture population inference. For topology inference, Also GTmix with small data is often more accurate in both the topology inference and the admixture population inference than TreeMix and MixMapper with large data in most cases. Note that TreeMix appears to outperform MixMapper when the number of populations increases.

**Figure 2:**
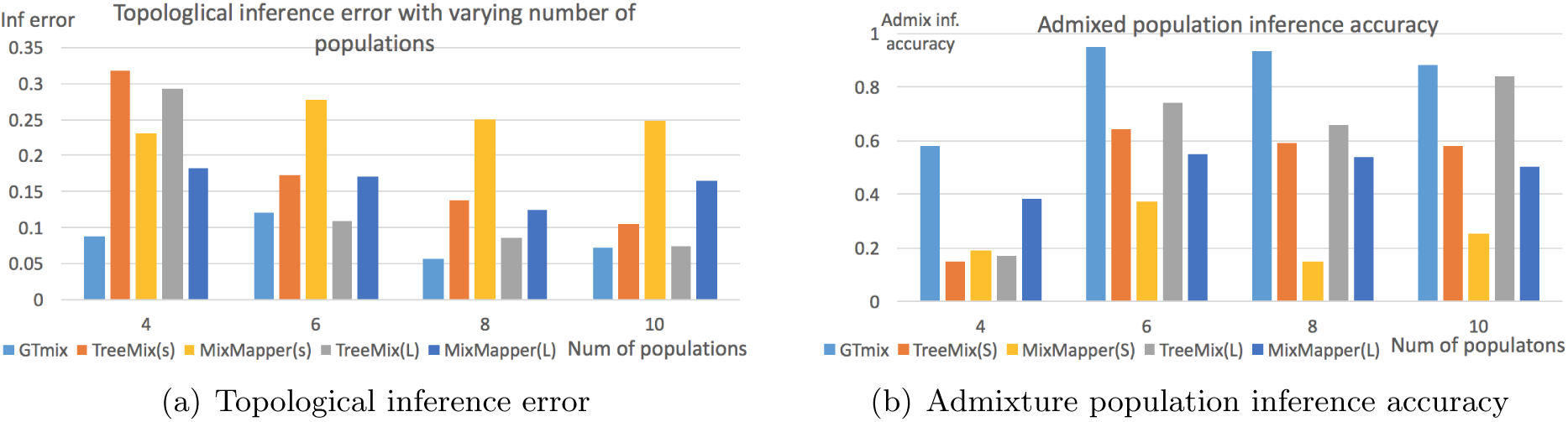
Admixture network inference with varying number of populations. GTmix: four alleles per population. TreeMix and MixMapper: small data (four alleles per population, denoted as “S”) and large data (100 alleles per population, denoted as “L”). Part 2(a): average best-match RF distance between inferred networks and true networks. Part 2(b): average admixture population inference accuracy. X-axis: number of populations.

##### Varying number of alleles per population

We evaluate the performance of GTmix when the number of alleles per population varies. Due to the computational difficulty, GTmix only runs on data with up to 8 alleles per population. For comparison, TreeMix and MixMapper are run on data with up to 100 alleles per population. Six populations are simulated. Figure 3(a) shows the results. As expected, when the number of alleles increases, all methods tend to be more accurate. On the same data, GTmix is consistently better than TreeMix and MixMapper. GTmix can be more accurate with smaller data than TreeMix and MixMapper even the latter are run with larger data. For example, the error of GTmix with 8 alleles per population is about 0.077, while the errors are about 0.11 and 0.17 respectively for TreeMix and MixMapper with 100 alleles per population.

**Figure 3:**
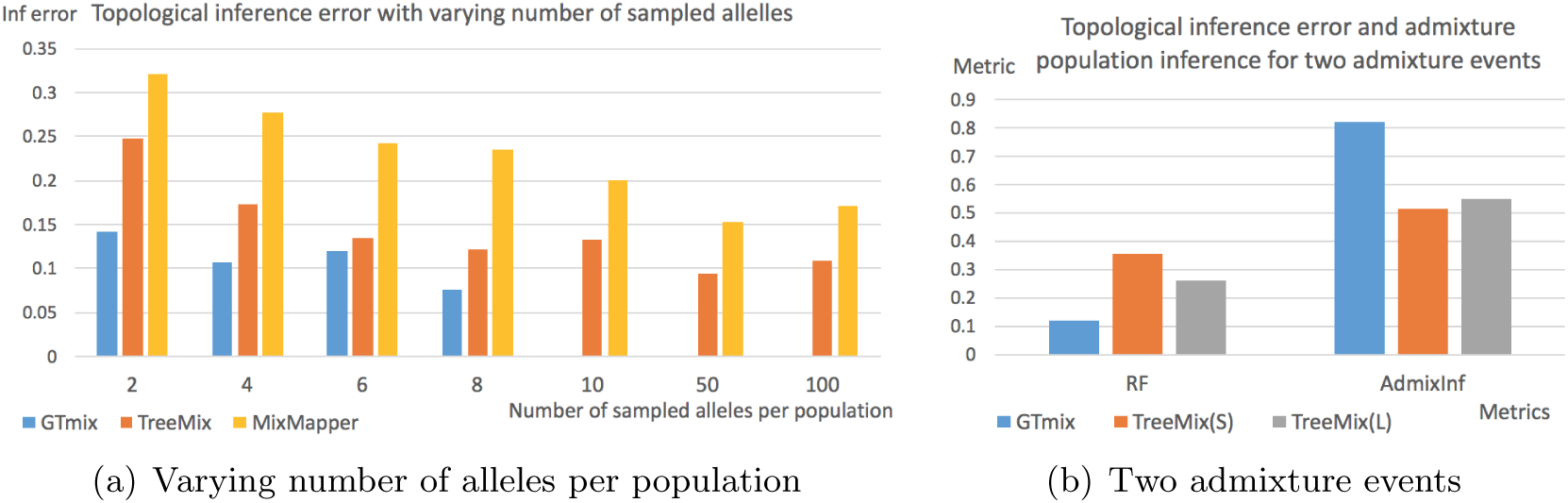
Network inference with varying number of alleles and number of admixture events. Part 3(a): varying number of sampled alleles per population. GTmix: 2 to 8 alleles per population. That is, GTmix is not run for 10, 50 and 100 alleles. TreeMix and MixMapper: 2 to 100 alleles per population. X-axis: number of alleles per populations. Part 3(b): with two admixture events; inference error for topology and admixture population inference accuracy. Six populations. TreeMix is run with both small (S) and large (L) data.

##### Two admixture events

By default, we simulate a single admixture event. To test the performance of GTmix on more complex demographic models, we simulate two admixture events for six populations (plus one outgroup). We compare GTmix (on small data) with TreeMix which are run on both small and large data. The results are shown in Figure 3(b). We can see that GTmix outperforms TreeMix in both topological inference and admixture population inference (on both small and large data) significantly even when the latter is run with large data.

##### Phasing

So far we use true simulated haplotypes. In practice, haplotypes are usually inferred from genotypes. We now evaluate the performance of GTmix on inferred haplotypes. Note that phasing accuracy can be affected by recombination rate. We simulate data with varying recombination rates (with default mutation parameter of 50). We randomly pair up two simulated haplotypes to form a genotype. Then we use the program beagle [1] to phase the gentoypes to obtain phased haplotypes. We compare the topological inference error of networks inferred with both the true haplotypes and the phased haplotypes.

The results are shown in Figure 4. When recombination rate is low, phasing error appears to have marginal effect on the inference accuracy. When recombination rate is high (say 250), the impact of recombination rate becomes larger: there is a significant reduction in inference accuracy when recombination rate is high.

**Figure 4:**
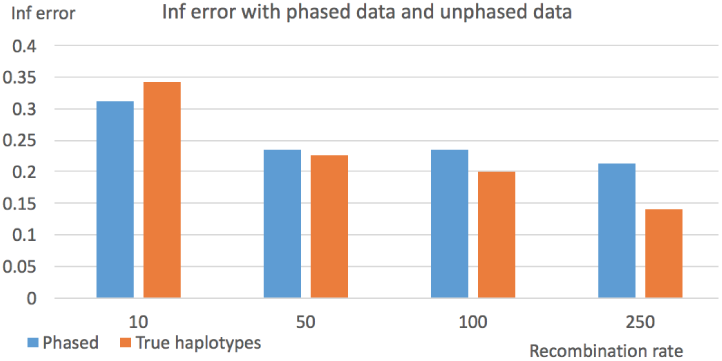
Topological inference error on phased and true haplotypes. X-axis: recombination rate used in data simulation.

#### 4.1.3 Network inference efficiency

We show the running time of GTmix under various settings in Figure 5. The running time of GTmix apparently grows exponentially with regard to both the number of populations and the number of alleles per population. On the other hand, GTmix scales well with regard to the number of loci. Overall, although GTmix is still computationally intensive on relatively large data, GTmix can be practical for inference for many data of current interests.

**Figure 5:**
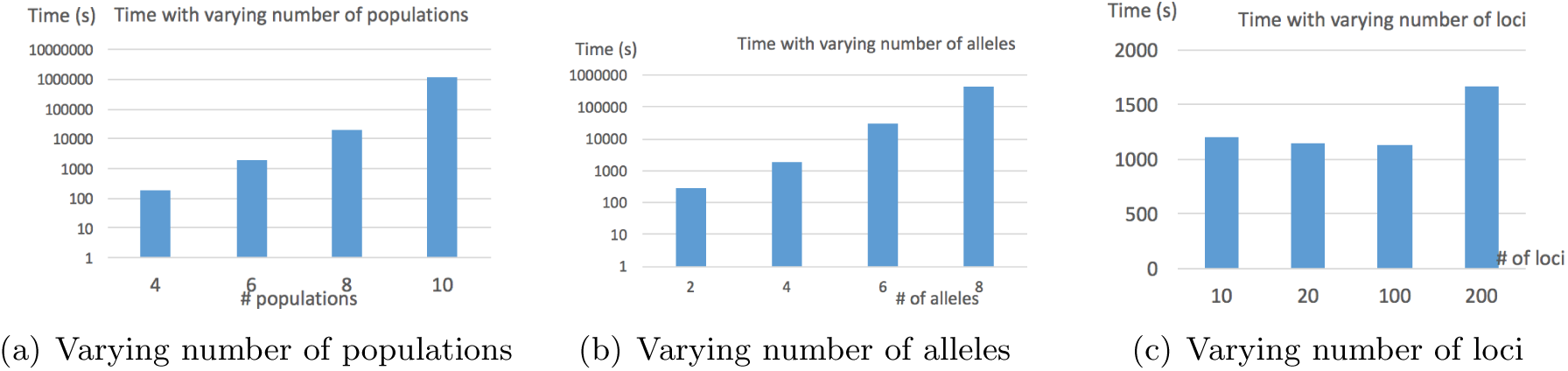
Running time of GTmix with varying number of populations, number of alleles per population, and number of loci. Default: six populations and four alleles per population.

### 4.2 Results on the 1000 Genomes Project data

The 1000 Genomes Project [3] has released haplotypes of 1,092 individuals from 26 populations in Phase I integrated variant set release. The 1000 Genomes Project defines five super populations: African, East Asian, European, South Asian and Admixed American. We pick two populations from each of these super populations. Namely, we pick the following ten populations: Americans of African Ancestry in SW USA (ASW), Yoruba in Ibadan, Nigeria (YRI), Han Chinese in Beijing, China (CHB), Japanese in Tokyo, Japan (JPT), Utah Residents with Northern and Western European Ancestry (CEU), Iberian Population in Spain (IBS), Gujarati Indian from Houston, Texas (GIH), Indian Telugu from the UK (ITU), Mexican Ancestry from Los Angeles USA (MXL) and Puerto Ricans from Puerto Rico (PUR). We create two test datasets, one small and one large. The small data contains haplotypes from two randomly chosen diploid individuals (i.e., four alleles) from each population. We sample up to 50 loci from chromosomes 1 to 10 as follows: for each chromosome, randomly sample up to 50 regions of 100 Kb each, which are evenly located as 3 Mb apart. (ii) The large dataset contains SNPs from twenty diploid individuals (i.e., ten times as many as the small data) from the *whole* genome (i.e., all SNPs are used). To reduce noise, any SNP sites with non-binary alleles are discarded.

We run GTmix to infer admixture network with two admixture events on the small dataset only. It takes about 34 hours for GTmix to find the optimal network in a computer cluster (with 2.1 Ghz CPU). The network is shown in Fig. 6(a). For comparison, we run TreeMix with whole genome data (with 20 individuals per population). We use the whole genome data here because TreeMix can run with large data. We perform preprocessing to discard rare variants: only SNPs with minor allele frequencies of 5% or larger are kept ^2^. This leaves total 7.63 million SNPs in the data. Note that this is about 50 times more data than what is used by GTmix. The resulting network is shown in Fig. 6(b). We note that the networks inferred by GTmix and TreeMix share some key topological properties. For example, both networks have MXL and PUR as admixed. On the other hand, the two networks also differ in some aspects. The network by GTmix, as expected, has IBS to be closely related to the source populations involved in both inferred admixture, while the other ancestral populations involved are closely related to CHB/JPT. While the true admixture network for these populations is not known, at least some aspects of the TreeMix’s network don’t agree with commonly accepted human demographic history. First, the rooting of the TreeMix network doesn’t agree with the “out-of-Africa” scenario. Also, an ancestral Eastern Asian population (which is after the divergence of CHB and JPT) is involved in the admixture of MXL but not for PUR. So potentially more accurate demographic history for these aspects are given in the network by GTmix, even when TreeMix uses 50 times more genetic data than GTmix. To summarize, our results show that GTmix can provide accurate inference of admixture networks on real data.

**Figure 6:**
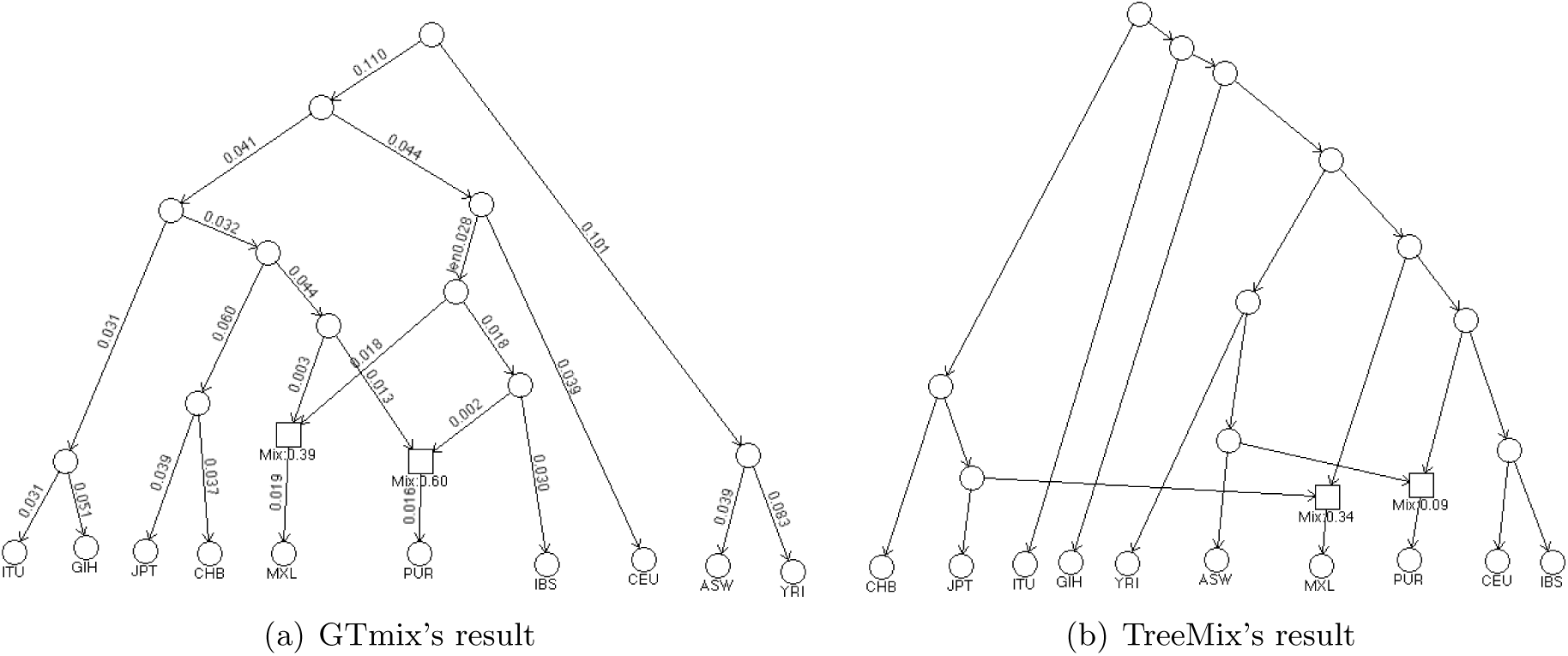
Inferred admixture network for ten populations from the 1000 Genomes Project by GTmix and TreeMix. GTmix is run with small data with two diploid individuals per population, while TreeMix is run with whole genome data with twenty diploid individuals per population (more than 50 times more data than that used by GTmix). Square box: admixed nodes (with inferred admixture proportions; the displayed admixture proportion is for the left source population). Each branch in the GTmix result has the estimated branch length (in the standard coalescent unit).

#### Different runs of GTmix and TreeMix

We have conducted more simulations with the 1000 Genomes data for GTmix and TreeMix. The purpose is testing the reproducibility and and the effect of data amount. Due to the space limit, the results are given in the Supplemental Materials.

#### Discussions

Due to the lack of space, discussion section is given in the Supplemental Materials.

## Software availability

The program GTmix can be downloaded from https://github.com/yufengwudcs/GTmix.

## ACKNOWLEDGMENTS

This work is partly supported by U.S. National Science Foundation grants CCF-1718093 and IIS1909425.

## Footnotes

1 There are over 34 million rooted binary trees with ten taxa. Adding two admixture nodes will lead to at least 3.4 × 10^11^ distinct networks.

2 If rare alleles are kept, our simulation results show that the constructed network appears to be less accurate.

