## Supplemental Materials for "Inference of Population Admixture Network from Local Gene Genealogies: a Coalescent-based Maximum Likelihood Approach"

#### S1 Additional background

##### S1.1 Gene genealogy

Gene genealogy $T$ of a set of sampled alleles from multiple populations can be very useful for the inference of population demographic history. For example, if alleles from populations $A$ and $B$ are closely located in $T$, then $A$ and $B$ may be closely related. On the high level, gene genealogy can be inferred from population haplotypes in a way similar to phylogeny inference. However, one major challenge is recombination. With recombination, gene genealogy of the sampled haplotypes is no longer a tree. Rather, a genealogy should be modeled by a genealogical network, called ancestral recombination graph (ARG) [6]. Note that it is often convenient to focus on local genealogical trees that are embedded in the ARG because trees are easier to work with than complex networks [28, 15]. For example, two local gene genealogies are shown in Figure 1(d) and Figure 1(e). Mutations can occur at a gene genealogy, which correspond to the single nucleotide polymorphisms (SNPs). A commonly used genetic model is the infinite sites model [10, 25], which implies that there is at most one mutation for each site in the sample history. Each sample allele has a haplotype at these SNP sites. See Figure 1(f) for an illustration of haplotypes. Haplotypes of sampled alleles offer hints on the underlying genealogy of these alleles. For example, in Fig. 1(f), haplotypes of $d_1$ and $d_2$ are identical, which suggests $d_1$ and $d_2$ are closely related in the genealogies. Inference of local genealogies from genetic data is a long standing difficult problem in population genetics. Recently, several new approaches [15, 24, 9] have been developed to infer genealogies from hundreds or even thousands haplotypes. A common feature of these tools is that they infer local genealogical trees. That is, these tools infer a list of genealogical trees from the given haplotypes, where each tree is for a specific site.

##### S1.2 Multispecies coalescent

Now suppose a genealogy $T$ has been inferred from the haplotypes. We start with a simple scenario: $\mathcal{N}$ contains a single population tree $\mathcal{T}_p$. Now for a fixed $\mathcal{T}_p$, the probability of observing a given genealogy $T$ assuming the underlying population tree is $\mathcal{T}_p$ is determined by the multispecies coalescent (MSC) model [23]. The MSC model assumes the independent coalescent process for each branch of $\mathcal{T}_p$. Due to MSC, there can be multiple different gene genealogies for $\mathcal{T}_p$ because of the stochasticity of coalescent process. To see this, look at the gene genealogy in Figure 1(d). Note that the lineage $b_2$ coalesces with the lineage $c_1$, even though $b_2$ and $c_1$ are from different populations (namely populations $B$ and $C$). Note that $b_2$ doesn't first coalesce with $b_1$ (which is from the same population $B$). This happens because by chance, $b_1$ and $b_2$ don't coalesce along the branch leading to the population $B$; when $c_1$, $b_1$ and $b_2$ meet at the parent of $C$, all three lineages can coalesce. By chance, $b_2$ coalesces with $c_1$ instead of $b_1$. This phenomenon is called incomplete lineage sorting (ILS), which is believed to be a common evolutionary process [23]. The MSC model offers a natural foundation to address the stochasticity in the coalescent.

One of the most important problems related to the MSC model is computing the probability of $T$ under $\mathcal{T}_p$. In the literature, this probability is called the gene tree probability. There are several algorithms for computing gene tree probability [4, 29, 31, 19]. Degnan and Salter [4] developed the first algorithm for computing gene tree probability. But their algorithm is very slow. We developed an algorithm, called the STELLS algorithm, [29], which is significantly faster than Degnan and Salter's algorithm. However, the STELLS algorithm and also another related algorithm [31] are still slow for large-scale inference. More recently, we developed a new algorithm called STELLS2 [19]. The STELLS2 algorithm is a heuristic. It computes an approximation of the gene tree probability and runs much faster than the algorithms in [29, 31]. Simulations show that the approximate gene tree probability computed by STELLS2 retains important information of the exact gene tree probability about the underlying population history [19]. In this paper, we adopt the STELLS2 algorithm for computing the (approximate) gene tree probability.

### S2 Additional methods

#### S2.1 Choosing a subset of genealogies for inference

##### S2.1.1 Choosing genealogies from each locus

GTmix uses the following simple method called TreePicker for picking a subset of gene genealogies from the (often large number of) inferred trees by RENT+. TreePicker works with haplotypes $\mathcal{H}_i$ from the $i$-th locus and the list of genealogical trees $\mathcal{T}_i$ inferred by RENT+ from $\mathcal{H}_i$. TreePicker chooses a fixed number $n_T$ trees from $\mathcal{T}_i$ as follows.

1. TreePicker first divides the SNPs within the $i$-th locus into $n_T$ equal-sized segments.
2. For each segment, TreePicker chooses one genealogy $T_i \in \mathcal{T}_i$ which "matches" the largest number of SNP sites within this segment.

Recall that a SNP site implies a split (bipartition) of sampled haplotypes under the infinite sites model. We say a tree $T$ matches a SNP site if there is a clade of $T$ whose leaves are exactly those on one side of the split implied by this SNP site. Intuitively, an inferred genealogy $T$ tends to be more reliable if it matches more SNP sites.

##### S2.1.2 Choosing a subset of trees to use for inference

GTmix takes a list of input trees chosen by TreePicker as input. The number of input trees can still be large, and these genealogies can still be noisy. Before GTmix starts the inference with inferred genealogies, it first chooses a subset of up to $K$ trees (by default, $K = 500$) from the input trees. The basic idea is removing input trees that are significantly different from the rest of trees. This is because such trees are more likely to contain errors. GTmix takes the following simple approach for choosing trees for inference. It first analyzes all the input trees and calculates the frequencies of the clades in the trees. Then, it scores a tree by clade frequencies: a tree $T$ is assigned a score $\prod_{C \in Clades(T)} freq(C)$. Here, $Clades(T)$ is the set of all clades of the tree $T$, and $freq(C)$ is the estimated frequency of the clade $C$ in all trees. GTmix chooses up to $K$ trees with highest scores.

#### S2.2 Initial population tree construction

Algorithm 2 is for constructing the initial population tree from a set of genealogies.

#### S2.3 Simulation

To test the performance of GTmix, we simulate haplotypes with regard to simulated admixture networks. There are a number of parameters in the simulation, which are shown in Table S1. Each parameter can have multiple values. Note that the number of combinations of all possible values over these parameters is very large. Thus, each parameter has a default value.

**Algorithm 2** Constructing initial population tree $\mathcal{N}_0$ from gene genealogies $\mathcal{T}$

```
Initialize D[p_i, p_j] to be 0 for all pairs of populations p_i and p_j.
for each T ∈ 𝒯 do
    for each pair of populations p_i and p_j with alleles in T do
        Let Lv(p_i) and Lv(p_j) be the set of leaves in T from p_i and p_j respectively.
        Let d_{p_i,p_j}(T, v_i, v_j) be the number of edges on the path from a leaf v_i ∈ Lv(p_i) to a
        leaf v_j ∈ Lv(p_j). Define d_{p_i,p_j}(T) to be the distance of populations p_i and p_j on T:
        d_{p_i,p_j}(T) is the the average of d_{p_i,p_j}(T, v_i, v_j) over all pairs of leaves v_i ∈ Lv(P_i) and
        v_j ∈ Lv(P_j).
        D[p_i, p_j] → D[p_i, p_j] + d_{p_i,p_j}(T)/|𝒯|
    end for
end for
Run neighbor joining on the distance matrix D to construct the initial population tree N_0.
```

##### S2.3.1 Admixture network simulation

We first generate random admixture networks with $n_p$ populations. Here, $n_p = 4, 6, 8,$ and 10. For each $n_p$, we simulate 10 randomly generated networks. We add one additional population as the outgroup. That is, the total number of populations is $n_p + 1$. A network has $n_a$ admixture nodes, where $n_a = 1$ (default) or 2. We focus on networks with more recent admixture in the simulation[3]. Except for a few networks with small number of populations (i.e., $n_p = 4$), we only choose extant populations (i.e., those at leaves of a network $\mathcal{N}$) as admixed populations. We arbitrarily order internal nodes in the simulated network $\mathcal{N}$, subject to the topological constraints imposed by the network. Then, each internal node $v_i$ (i.e., the $i$-th node in the order) is assigned a time $t_i = i \times t_s$ where $t_s$ is the time interval between events. All leaf nodes have zero time. The outgroup is assigned a relatively large time $t_o$, which by default is 0.3. The default value of $t_s$ is 0.02 in the standard coalescent unit. Each admixture node $v$ is associated with an admixture proportion $m_v$. By default, $m_v = 0.5$. See the Supplemental Materials for the list of simulated networks with four and six populations.

##### S2.3.2 Haplotype simulation

For each simulated admixture network $\mathcal{N}$, we simulate a set of haplotypes using the program ms [7]. For each population, we simulate $n_c$ alleles. By default, $n_c = 4$. We simulate $n_L$ loci for each setting. By default, $n_L = 500$. For each locus, we set the mutation parameter $\mu$ (by default, $\mu = 50$), and the recombination parameter $\rho$ (by default, $\rho = 50$) with region length being $500,000$ base pairs. The island model with population admixture implemented in ms is used to simulate the haplotypes on a specific admixture network.

#### S3 Networks used in simulation

The ten randomly chosen networks used in simulation are shown in Figure S1 (for four populations) and Figure S2 (for six populations). Note that each network has one more population as the outgroup (not shown).

[3] Simulations show that ancestral admixture is more difficult to infer for all inference schemes.

Table S1: *A list of parameters and their default values used in the simulation.*

| Description | Symbol | Default |
|---|---|---|
| Number of populations (excluding the outgroup) | $n_p$ | 6 |
| Number of admixture nodes in a network | $n_a$ | 1 |
| Time interval between events (in coalescent units) | $t_s$ | 0.02 |
| Admixture proportion at admixture node $v$ | $m_v$ | 0.5 |
| Number of alleles per population | $n_c$ | 4 |
| Number of loci | $n_L$ | 500 |
| Mutation parameter | $\mu$ | 50 |
| Recombination parameter | $\rho$ | 50 |
| Length of locus | $L$ | $500,000$ |

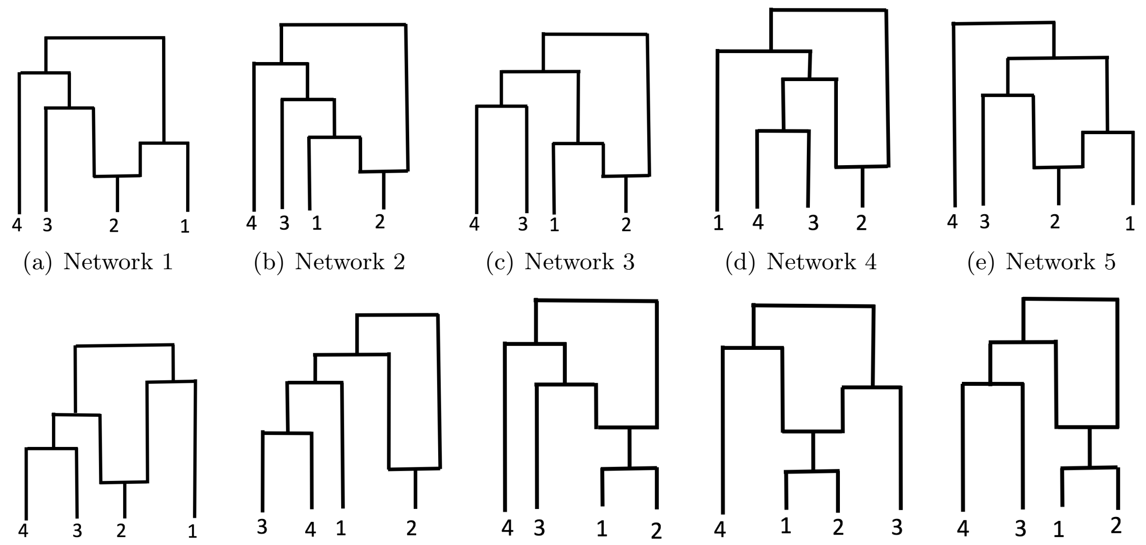


(a) Network 1 (b) Network 2 (c) Network 3 (d) Network 4 (e) Network 5

(f) Network 6 (g) Network 7 (h) Network 8 (i) Network 9 (j) Network 10

Figure S1: *Simulated admixture networks with four populations. Each network has one additional population as the outgroup (not shown). Note that the admixture events in parts 1(h) to 1(j) occur at ancestral populations, not extant populations.*

### S4 Additional Results

#### S4.1 Network inference with different admixture proportions

The default admixture proportions used in simulation is 0.5 (i.e., equal contribution from each parental populations at admixture). It is interesting to investigate how GTmix and TreeMix perform with different admixture proportions. We simulate haplotypes with varying admixture proportions: $0.1, 0.3, 0.5, 0.7,$ and $0.9$. Six populations are simulated. All the other parameters are in their default values. The average network inference topology error (i.e., the best-match RF distance) are shown in Figure S3. It can be seen that GTmix outperforms TreeMix on small data on all admixture proportions. GTmix also has similar network inference accuracy as TreeMix which is run on large data in different admixture proportions. We also note that all methods perform better for admixture proportions near 0.5 than those far from 0.5.

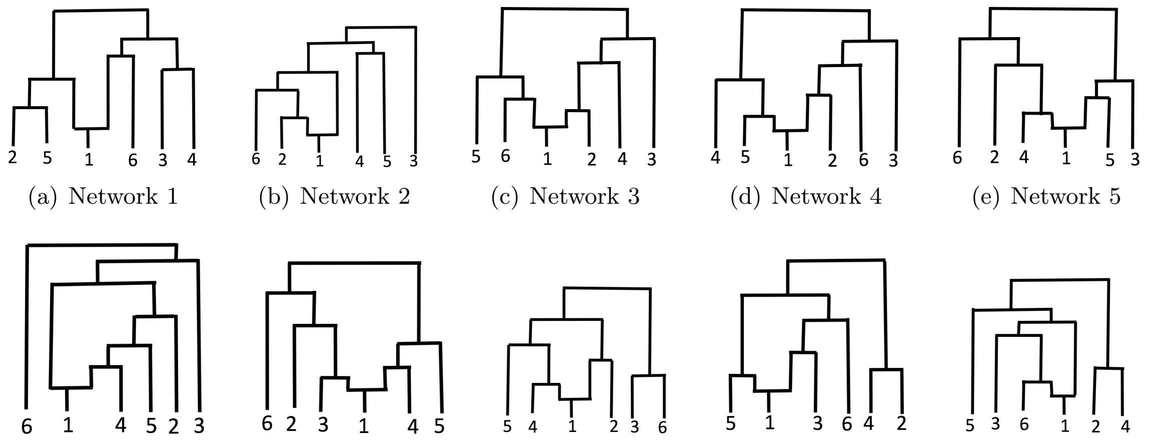


(a) Network 1 (b) Network 2 (c) Network 3 (d) Network 4 (e) Network 5

(f) Network 6 (g) Network 7 (h) Network 8 (i) Network 9 (j) Network 10

Figure S2: *Simulated admixture networks with six populations. Each network has one additional population as the outgroup (not shown).*

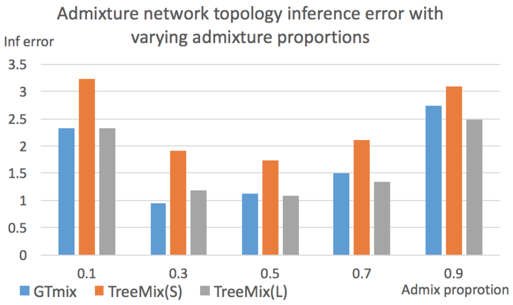


Figure S3: *Topological inference error on data with varying admixture proportions. Six populations. X-axis: admixture proportions. Y-axis: topological inference error (the best-match RF distance). GTmix is run with small data (four alleles per population). TreeMix is run with both small (denoted as S) and large (denoted as L, with 100 alleles per population).*

#### S4.2 Number of loci, recombination rate and mutation rate

There are several additional factors that may affect the inference accuracy. We now evaluate the influence of the number of loci, recombination rate and mutation rate on admixture network inference. We simulate data with various number of loci, recombination and mutation rates. We show both topological inference error and admixture population inference accuracy. Results are shown in Figure S4. As expected, when the number of loci increases, network inference becomes more accurate. The same occurs when the mutation rate increases. This is because when mutation rate increases, the number of SNPs increases and the inferred gene genealogies become more accurate (see, e.g., [15]). For recombination rate, it is a little surprising that increasing recombination rate can make the network inference more accurate. Note that when recombination rate increases, it becomes more difficult to infer gene trees [15]. Thus, it appears that moderate decrease in the accuracy of inferred genealogies doesn't have a major impact on the admixture network inference.

#### S4.3 Admixture proportion inference

In addition to topology and admixture population inference, it is also desirable to infer the admixture proportions. We now run GTmix and TreeMix on data with various admixture proportions. Here we simulate four populations. We report the average absolute error between the inferred

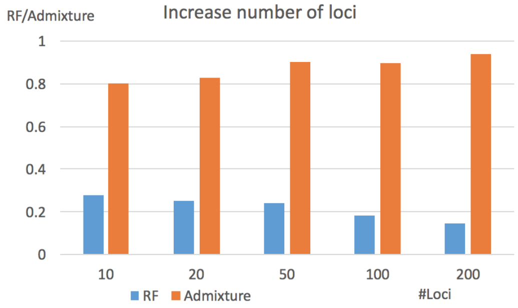


(a) Varying number of loci

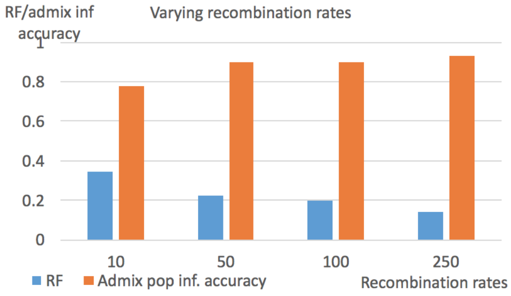


(b) Varying recombination rate

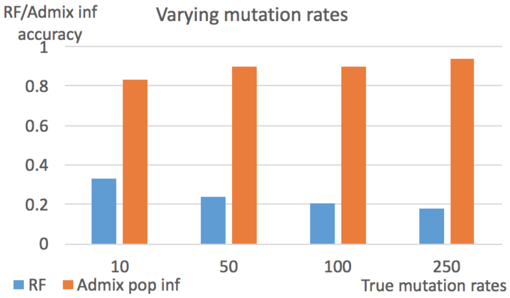


(c) Varying mutation rate

Figure S4: *Performance of admixture network inference with varying number of loci, recombination rate and mutation rate. Part 4(a): varying number of loci. Part 4(b): varying recombination rate. Part 4(c): varying mutation rate. X-axis: parameter setting. Y-axis: RF error/admixture population accuracy.*

proportion and the true proportion. GTmix is run on small data, while TreeMix is run on small data and large data. The results are shown in Figure S5. We can see that GTmix outperforms TreeMix on small data. Compared with TreeMix with large data, GTmix with small data performs well when the true proportions are near 1/2, and is less accurate than TreeMix with large data when the true proportions are close to 0 or 1.

#### S4.4 Varying the number of genealogies for inference

The running time in Fig. 5(c) doesn't grow linearly with regard to the number of loci. The main reason is that GTmix samples a fixed number $K$ of trees from the given set of trees. Here, the default value of $K$ is fixed to be 500. The choice of $K$ can affect both the running time and also accuracy of GTmix. To investigate the effect of the value of $K$, we test GTmix with varying $K$ values (from 100 to $5,000$) with six populations and four alleles per population. Fig. S6 shows the results. There is a clear trade-off between inference accuracy and efficiency with regard to the number of sampled trees. Overall, larger $K$ values tend to produce more accurate inference results, although the running time will be larger. In practice, therefore, it may be beneficial to use larger $K$ when the running time remains acceptable.

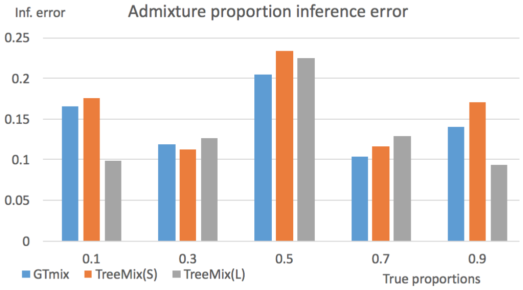


Figure S5: *Admixture proportion inference for GTmix on small data, TreeMix on small data (denoted as S) and TreeMix on large data (denoted as L). X-axis: true admixture proportions. Y-axis: absolute admixture proportion inference error. Four populations.*

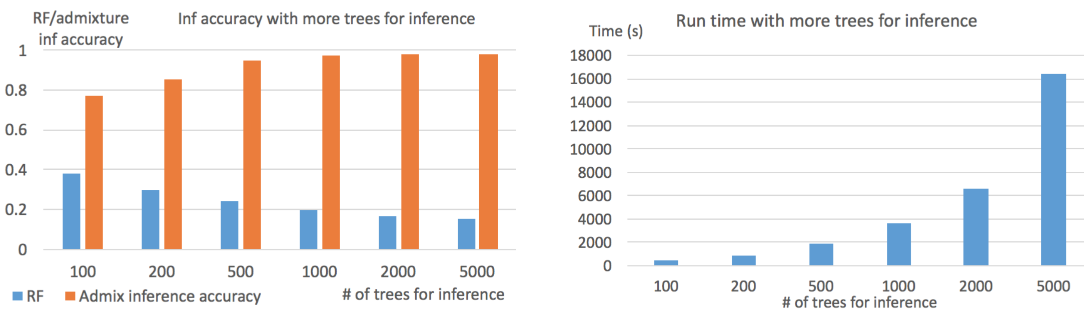


(a) Inference error

(b) Time (s)

Figure S6: *Effect of the number of sampled trees for inference on accuracy and running time. Part 6(a): network topology inference error (best-match RF distance) with varying number of sampled trees $K$. Part 6(b): running time (in seconds) with varying $K$.*

##### S4.5 More results on the 1000 Genomes Data

**Different runs of GTmix** To test how reproducible the results by GTmix are, we run GTmix for four more times using disjoint sets of sampled individuals from each population. We use the same procedure as described above. Among these four independent tests, GTmix reports MXL and PUR as admixed in three out of four tests. Moreover, the inferred admixture networks in these three tests are similar to those reported by the original test (with some differences in the location of the source populations for some admixture events). This indicates that GTmix is overall consistent in its inference, although there is some variance in the results when different individuals in the populations are used for inference.

**Different runs of TreeMix** We now evaluate the effect of data amount on TreeMix inference. We first downsample the whole genome data by randomly picking one fifth SNPs for total 200 diploid individuals (20 individuals per population). There are about 1.5 million SNPs left (which is still about 10 times as large as those used by GTmix). Here, we use the same 20 individuals from each population as before. Thus, even with downsampling, TreeMix is run with data that is about 100 times larger than that is used for GTmix. Here, rare SNPs are discarded: only SNPs with minor allele frequencies of 5% or larger are kept [4]. We repeat the downsampling four times. The inferred networks are shown in Figure S7. While MXL and PUR are consistently admixed in these networks, the population trees among these runs are significantly different from each other (especially the rooting).

#### S5 Discussions

Inference of admixture network from genetic data is challenging computationally. A common approach is considering each SNP site independently. Single site based inference methods can be based on moments [20, 13] or likelihood based (see, e.g., [2]). While these methods are usually computationally faster than full likelihood methods, a major downside is that these methods don't consider linkage equilibrium (LD) that is captured by population haplotypes. The effect of LD on local genealogies is that nearby local genealogies tend to be highly similar [28]. Note that most phylogeny inference methods work with sequences, not individual variation sites. This is because sequences capture more information about the underlying evolutionary process than individual sites. Thus, admixture network inference using haplotypes, rather than individual SNPs, may improve the accuracy. As shown in this paper, this is indeed the case. Haplotypes can be more difficult to process than single variation sites. A key point made by this paper is that handling

[4] If rare alleles are kept, our simulation results show that the constructed network appears to contain more errors.

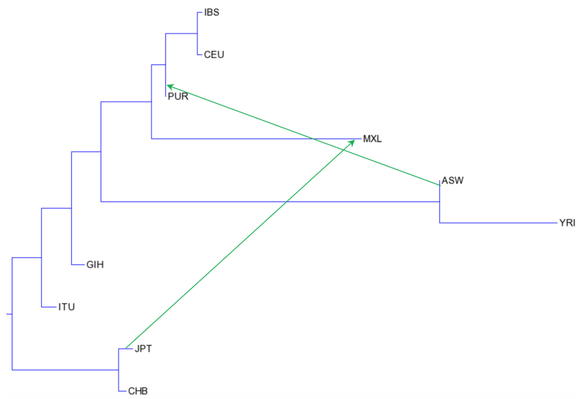


(a) Network with 20% downsampling, test 1

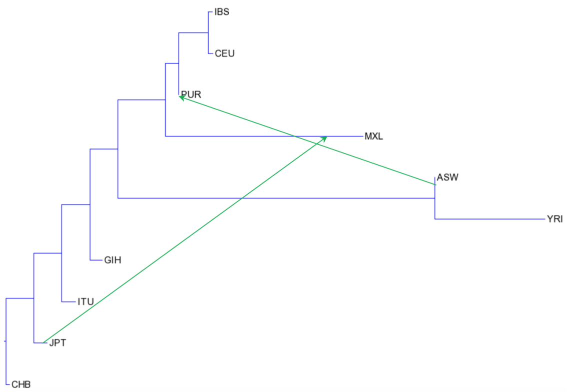


(b) Network with 20% downsampling, test 2

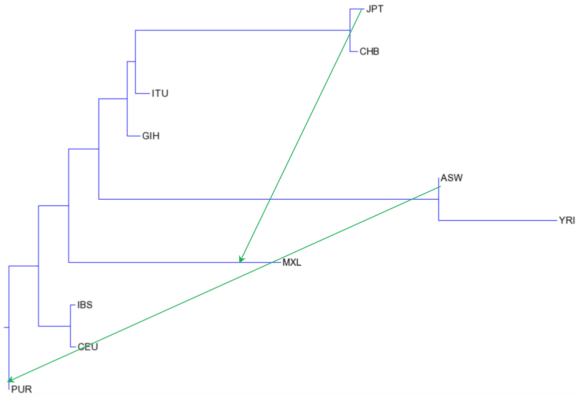


(c) Network with 20% downsampling, test 3

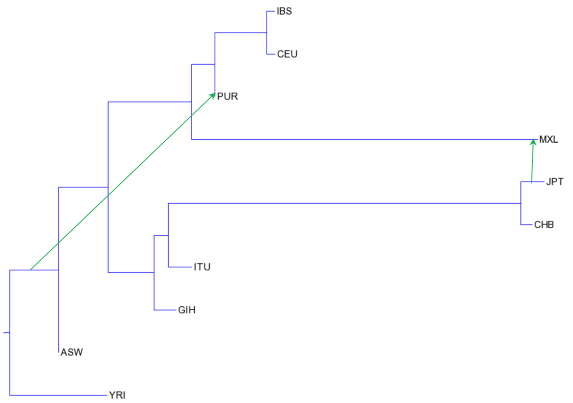


(d) Network with 20% downsampling, test 4

Figure S7: *Admixture network for ten populations inferred by TreeMix using whole genome data, after discarding rare SNPs (with 5% or less minor allele frequencies). Parts 7(a) to 7(d): four 20% downsampling tests. Admixture proportions (not shown): similar to the original inference. Branch length: roughly proportional to the inferred lengths by TreeMix.*

larger amount of data doesn't necessarily lead to more accurate results. A slower network inference method can potentially provide more accurate results even with smaller data, if this method can utilize the given data more effectively than existing methods. We show in this paper that our new method, GTmix, utilizes the given genetic data more effectively than existing methods.

Ideally, we would like to compute the likelihood of haplotypes for a given admixture network. However, it is computationally difficult to compute such likelihood. In contrast, the likelihood of gene genealogies is known to have (relatively) efficient algorithms under certain conditions. The key idea of GTmix is relying on the likelihood of inferred gene genealogies. This gets around one of the major computational challenges for coalescent likelihood based inference of admixture network. Local genealogy is not easy to infer due to recombination. There is still significant noise in the inferred local genealogies by any existing genealogy inference methods. It is conceivable that the noise in the inferred genealogies is so large that outweighs any benefits of using the inferred genealogies. In this paper, we show that this appears to not be the case for admixture network inference: GTmix can produce more accurate inference results using inferred genealogies than methods not using genealogies. Our results suggest that LD can indeed reveal important aspects about population demographic history. It has been shown [24, 9] that the inferred gene genealogies can be very informative for population genetic inference (e.g., for natural selection and demographic history). In this paper, we show that inferred gene genealogies can be very informative for admixture network inference.

GTmix is based on coalescent likelihood of inferred local genealogies. Inferring population admixture network by maximizing coalescent likelihood is a natural approach because the coalescent process captures the key aspects of population admixture. We note that coalescent likelihood based inference has been used (in software tools such as PhyloNet [26]) to infer phylogenetic network, which on the high level is related to admixture networks. However, inference of phylogenetic networks is mainly applied on species level data, which is quite different from population genetic data. Moreover, when coalescent likelihood is used, the PhyloNet approach is still quite limited in terms of the size of phylogenetic networks [32] it can work with[5]. So far, we are not aware of any existing coalescent likelihood based admixture network inference methods that have been used widely by population geneticists. A main challenge is the computational difficulty for computing coalescent likelihood of an admixture network. Thus, it is commonly believed that coalescent likelihood is difficult to be applied on data of practical interests [20]. GTmix takes advantage of the recent progress toward efficient computation of gene tree based coalescent likelihood [29, 31, 19]. Now it becomes feasible to compute gene tree based coalescent likelihood for gene trees that are of practical interests. This enables GTmix to infer admixture networks with ten or more populations, which can be of current interests to geneticists. GTmix takes an approximation when calculating the gene tree probability for a network: instead of considering the full network, GTmix only considers the set of population trees in the network. This approximation appears to work reasonably well in our simulation. Nonetheless, computing coalescent likelihood of gene trees remains a challenging computational problem, even with the population tree approximation. At present, GTmix cannot handle large data (e.g., 20 populations with 10 sampled alleles per population). In contrast, existing methods such as TreeMix and MixMapper can handle much larger data. Much more work is still needed to further speed up the likelihood computation for more efficient inference.

Existing local genealogy inference tools can infer coalescent time. Note that coalescent time can carry information about population demographic history. GTmix only uses the inferred topology (i.e. it doesn't use coalescent time). The probability of a gene genealogy is the probability of the topology of the gene genealogy. This is because our experience indicates that there is often significant noise in the inferred coalescent time, which can lead to less accurate inference results [29].

There are two ways to speed up the network inference by GTmix. First, when the number of populations is large, one may choose a subset of populations to work with. Our experience indicates that GTmix usually is reasonably fast for up to ten populations. For large datasets, it may be useful to infer networks for different subsets of populations and then consider merging these networks. We admit, however, merging networks from different subsets of populations may not be very straightforward. This is because of inference errors of these networks. One should be aware of potential inference errors if such merging is attempted. Second, reducing the number of alleles may speed up the inference. Note, however, reducing the number of alleles can reduce the inference accuracy as well. There is a trade-off between accuracy and efficiency here.

In this paper, we focus on admixture network inference. We note that population demographic history is complex and admixture network only captures part of demographic history. There are several other important demographic aspects not addressed by GTmix. For example, it is widely known that population sizes can fluctuate (see, e.g., [12]). There can also be complex forms of gene flow that is not captured by the current admixture network formulation. Putting all aspects of demographic history in a single model and performing inference with the model is a daunting challenge computationally. Inferring more complete demographic population history from genetic data will need major progress in computational methodologies.

[5]PhyloNet implements different phylogenetic network inference methods; the MSC-based inference appears not be able to infer some networks constructed by GTmix (say ten taxa and two admixture nodes).
